# The metabolic profile of Extracellular Vesicles identifies and separates patients with Sarcoidosis and Anti-Synthetase Syndrome

**DOI:** 10.64898/2026.05.05.722727

**Authors:** Loïc Steiner, Maria Eldh, Christina Glykeria Samakovli, Elga Bernardo Bandeira De Melo, Hasnat Noor, Ralph Evaristo Claret Monte, Chantal Reinhardt, Carl Wengse, Maryam Fathi, Begum Horuluoglu, Anders Lindén, Lena Palmberg, Ingrid E. Lundberg, Susanna Kullberg, Gözde Güçlüler Akpinar, Susanne Gabrielsson

**Affiliations:** Division of Immunology and Respiratory Medicine, Department of Medicine Solna, Karolinska Institutet, Stockholm, Sweden; Center for Molecular Medicine, Karolinska University Hospital, Stockholm, Sweden; Clinical Immunology and Transfusion Medicine, Karolinska University Hospital, SE-171 77 Stockholm, Sweden; Institute of Environmental Medicine, Karolinska Institutet, SE-171 77 Stockholm, Sweden; Department of Internal Medicine and Infection, Danderyd Hospital, Danderyd, Sweden; Division of Rheumatology, Department of Medicine, Solna, Karolinska Institutet, Stockholm, Sweden; Department of Gastro, Dermatology and Rheumatology, Theme Inflammation and Aging, Karolinska University Hospital, Stockholm, Sweden; Department of Respiratory Medicine and Allergy, Theme Inflammation and Ageing, Karolinska University Hospital, SE-171 76 Stockholm, Sweden

**Author notes:** Shared senior authorship (Gözde Güçlüler Akpinar, Susanne Gabrielsson). **<u>Corresponding author</u>**: Susanne Gabrielsson, Division of Immunology and Respiratory Medicine, Dept. of Medicine, Karolinska Institutet Center of Molecular Medicine (CMM) L8:03 SE-171 64, Solna, Sweden.

## Abstract

Sarcoidosis is a multisystem disorder that primarily affects the lungs and is characterizedby granulomatous inflammation. However, much of the underlying disease mechanisms remain poorly understood. Extracellular vesicles (EVs) are small membrane-bound particles released by all cells and carry various cargos including metabolites. They are involved in intercellular communication that can be dysregulated in diseases.This study characterizes the metabolic cargo of EVs isolated from bronchoalveolar lavage fluid (BALF), using liquid chromatography-mass spectrometry (LC-MS)-based metabolomic analysis, in patients with sarcoidosis (n=37), compared to healthy controls (n=10). Additionally, the sarcoidosis signature was compared to another pulmonary disorder, anti-synthetase syndrome (ASyS, n=10). Arachidonic acid (AA) results were verified by ELISA.

A total of 1202 metabolites were detected, with 111 annotated ones further analyzed. EVs from sarcoidosis patients showed distinct metabolomic profiles compared to both ASyS patients and healthy controls, with 38 annotated metabolites differentially expressed in any of the groups. In both annotated and non-annotated data, sarcoidosis patients clustered separately from ASyS patients and healthy individuals. Furthermore, sarcoidosis patients clustered in 3 subgroups, whereof one was similar to ASyS patients and one stood out as showing higher cell counts in BALF. Higher AA levels were found in sarcoidosis patient EVs by LC-MS, and AA results were verified by ELISA.

Our data show that BALF EV metabolites are disease-dependent and support the notion thatsarcoidosis patients should be further subgrouped for better diagnosis and treatment.

## Introduction

Sarcoidosis is a complex systemic inflammatory disorder of unknown etiology. It is characterized by the formation of granulomas in different organs, mainly in the lungs. Pulmonary sarcoidosis leads to impaired breathing ability, decreased quality of life, and increased mortality.

Sarcoidosis can be present in different clinical phenotypes, and its progression varies among individuals. While some patients can recover without any treatment, particularly those with Löfgren’s syndrome (LS), another group of individuals (mainly patients with non-LS) develop chronic lung inflammation that may sometimes lead tofibrosis. It is believed that, in genetically predisposed individuals, exposure to unknown antigen(s) and/or other environmental factors leads to the development of the disease (1). The immunological hallmarks of sarcoidosis include activation of macrophages and accumulation of CD4^+^ T cells in affected organs, eventually leading to the formation of granulomas. However, the underlying mechanisms responsible for the initiation and persistence of inflammation are still poorly understood (2), and unpredictable treatment outcomes suggest unknown mechanisms and subgroups of the disease.

Anti-synthetase syndrome (ASyS) is an autoimmune disease characterized by the presence of autoantibodies targeting one of several aminoacyl t-RNA synthetases along with clinical features including interstitial lung disease, myositis, Raynaud’s phenomenon, arthritis, mechanic’s hands, and fever (3, 4). One of the most common autoantibodies in this disorder is anti-Jo1, targeting histidyl-transfer RNA synthase. Up to 90% of the patients with anti-Jo1 ASyS present lung manifestations with symptoms that overlap with those of sarcoidosis, which may make the distinction difficult at the time of diagnosis. Thus, more knowledge about the mechanisms, as well as new biomarkers that separate and categorize these diseases are needed.

Extracellular vesicles (EVs) are small, nanosized membrane-enclosed vesicles that mediate cellular communication. They encapsulate a vast cargo of bioactive molecules, such as nucleic acids, proteins, lipids, and metabolites, reflecting the status of their originating cells (5, 6). Additionally, their cargo can be transferred to recipient cells, modulating their physiological state (7). These features, combined with their abundance in diverse biological fluids, justify studying their contents to better understand disease mechanisms, potentially leading to their use as therapeutic targets or biomarkers.

The metabolic composition of EVs has recently gained interest (8), both as a novel way of modulating the metabolism of recipient cells and as new biomarkers are detected in the vesicles. Substantial evidence suggests that alterations in metabolic pathways may influence the inflammatory profile of diseases (9-11). To draw the metabolomic landscape of EVs in the lung and identify disease-specific signatures, we aimed to investigate and compare the metabolite cargo of EVs derived from bronchoalveolar lavage fluid (BALF) of patients with sarcoidosis and ASyS, as well as healthy controls. We found significant differences between the disorders studied and identified specific metabolites and pathways enriched in each condition, suggesting that there is a unique pathology reflected in the metabolome of EVs. We further report that the sarcoidosis patients divide into three distinct subgroups and higher levels of arachidonic acid (AA) were verified by ELISA. This first metabolome study of human BALF EVs aids in the understanding of the mechanisms of sarcoidosis and ASyS and highlights distinct EV-associated metabolic patterns that could form the basis for improved diagnostics and targeted interventions, leading to better quality of life for patients.

## Materials and Methods

### Patients

All patients and healthy controls gave informed consent for this study approved by the local ethics committee. Bronchoscopy to collect BAL was performed at Karolinska University Hospital, Sweden, for all patients as previously described (12). The bronchoscopy was used as part of routine diagnostics procedures (for sarcoidosis and ASyS patients), or as included in the study protocol for healthy controls. All diagnoses were validated by a respiratory physician and established according to international guidelines (13-16). Detailed patient characteristics are presented in Supplementary Table 1.

### Extracellular Vesicle isolation

EVs were isolated as described previously (7) in accordance with the MISEV guidelines (17). After BALF collection, EVs were enriched by differential ultracentrifugation (10 min at 400 x g, 30 min at 3,000 x g, 30 min at 10,000 x g, 0.22µm filtration, followed by a centrifugation and wash in PBS for 2h at 100,000 x g. All high-speed centrifugations were performed using a fixed-angle rotor (Ti45, Beckman-Coulter) at 4°C.

### EV characterization

In our previous publications, we thoroughly characterized EVs from the BALF of healthy controls and sarcoidosis patients (18, 19). These EVs were isolated using identical protocols and settings, and were analyzed for morphology, total protein amounts, full proteomic content, and non-vesicular contaminations (19). Here, basic characterizations of the EVs were made in a similar manner.EVs were quantified and analyzed for size using a Nanoparticle Tracking Analysis (NTA, LM10 platform with sCMOS camera from NanoSight Ltd., Salisbury, UK, 40nm laser, camera level 10, detection threshold 3).

Markers on the EV surface were detected by bead-based flow cytometry analysis, with anti-CD9-coated magnetic beads (Spherotech SVMS-40-10 with anti-CD9 antibody, HI9a, Biolegend cat#312112) and PE anti-CD9 detection antibodies (HI9a, Biolegend cat#312106). The stained EV-beads complexes were acquired on a FACSCanto II flow cytometer (BD). FCS files were further analyzed using FlowJo^TM^ 10 software (BD).

### Metabolite extraction and metabolomic analysis

EV samples, isolated from 10 mL of BALF from healthy controls, ASyS, and sarcoidosis patients, were sent to the Swedish Metabolomics Centre for LC-MS-based untargeted metabolomics analysis.

### Data processing

The metabolites detected by the untargeted approach were compared to a database of well-characterized metabolites curated by the Swedish Metabolomics Centre, yielding an annotated dataset with high identification confidence. The metabolites were filtered to keep the ones detected in at least 20% of the samples per group at an intensity above the blank. After log2 transformation, missing values were imputed (95% of the minimally detected value for that specific metabolite across each sample), and the metabolite intensities were median-normalized prior to further comparisons.

### Arachidonic acid enzyme-linked immunosorbent assay (ELISA)

Arachidonic acid (AA) levels were quantified using a commercially available competitive ELISA kit (Nordic Biosite, Täby, Sweden, Cat. No. EK2F239) according to the manufacturer’s instructions. In brief, EVs isolated from 29 sarcoidosis patients corresponding to 3 mL of original bronchoalveolar lavage fluid (BALF) were diluted to a final volume of 50 µL. Standards and samples were added to the pre-coated wells of the plate together with biotin-labeled detection antibody and incubated at 37 °C. After washing, HRP-streptavidin conjugate was added, followed by the 3,3′,5,5′-tetramethylbenzidine (TMB) substrate. The reaction was terminated by adding stop solution and absorbance was measured at 450 nm using the Varioskan™ LUX Multimode Microplate Reader. The standard curve was plotted in GraphPad prism (version 11.0.0). The samples’ AA concentrations were determined from the standard curve.

### Statistical analyses

The quantified LC/MS results were analyzed with R (version 4.2.2) in RStudio with open-source R packages including ‘metabolomics’, ‘dplyr’, and ‘pcaMethods’. Data was visualized using GraphPad prism (version 11.0.0). Enrichment analyses were done on the median normalized dataset using the ‘metaboAnalyst 6.0’ web software (20). The code does not contain any newly generated software, but it is available from the corresponding author upon request.

## Results

### Extracellular vesicles isolated from BALF show a typical small EVs phenotype

The BALF samples were collected from a cohort of 37 sarcoidosis patients, 10 patients with ASyS, and 10 healthy controls, as shown in Table 1 and Supplementary Table 1. EVs from the same volume of BALF were isolated and sent for metabolomic analysis (sample processing scheme in Figure 1A). Nanoparticle tracking analysis was used to measure the average particle size for each experimental group (Figure 1B), showing no differences in size between EVs from the two disease groups or healthy controls. Interestingly, bead-based flow cytometry revealed discrepancies in CD9 expression between groups, with a lower signal observed on EVs from healthy controls, compared to samples from patients with sarcoidosis and ASyS (Figure 1C).

**Figure 1:**
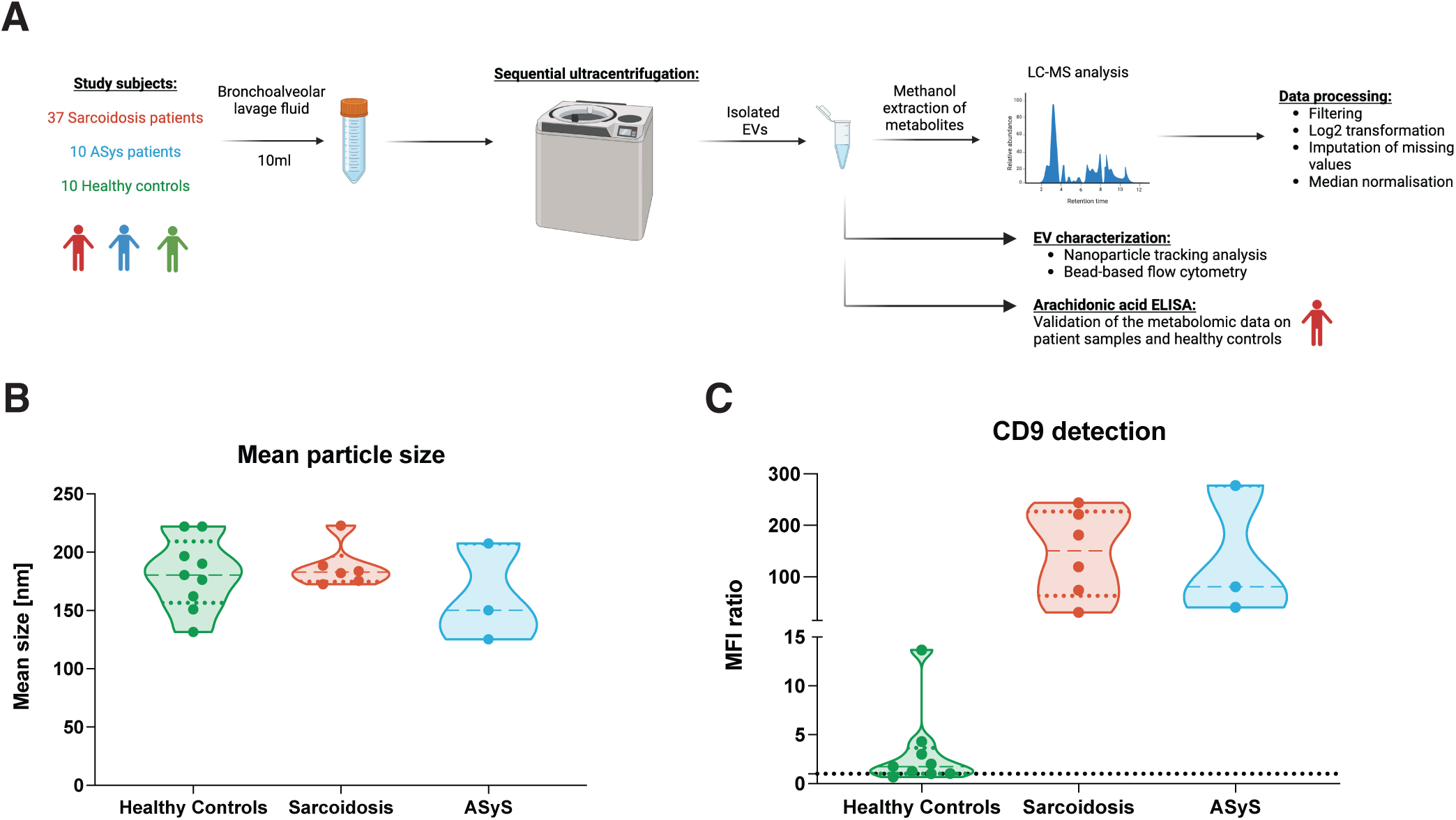
EVs isolated from BALF of healthy controls or patients with sarcoidosis and anti-synthetase syndrome (ASyS) show typical small EV characteristics. A. Schematic outline of the cohort, sample processing, analysis and validation. B. Nanoparticle tracking analysis showing the size distribution of samples originating from different groups. C. Bead-based flow cytometry of EVs captured with anti-CD9 coated beads and detected with anti-CD9 antibodies (MFI=mean fluorescence intensity, normalized to the isotype control, n=3-9 samples per group).

### The metabolic content of EVs shows a strong clustering of samples based on their disease of origin

A total of 1202 metabolites were initially detected in an untargeted way, and a subset of metabolites identified with high confidence was further annotated and considered for this study. When looking at the untargeted list of compounds in the samples, the heatmap of metabolite expressions revealed a clustering of samples based on the disease and clusters of metabolites were observed to be differentially expressed between the EVs from different lung disorders (Figure 2A and B).

**Figure 2:**
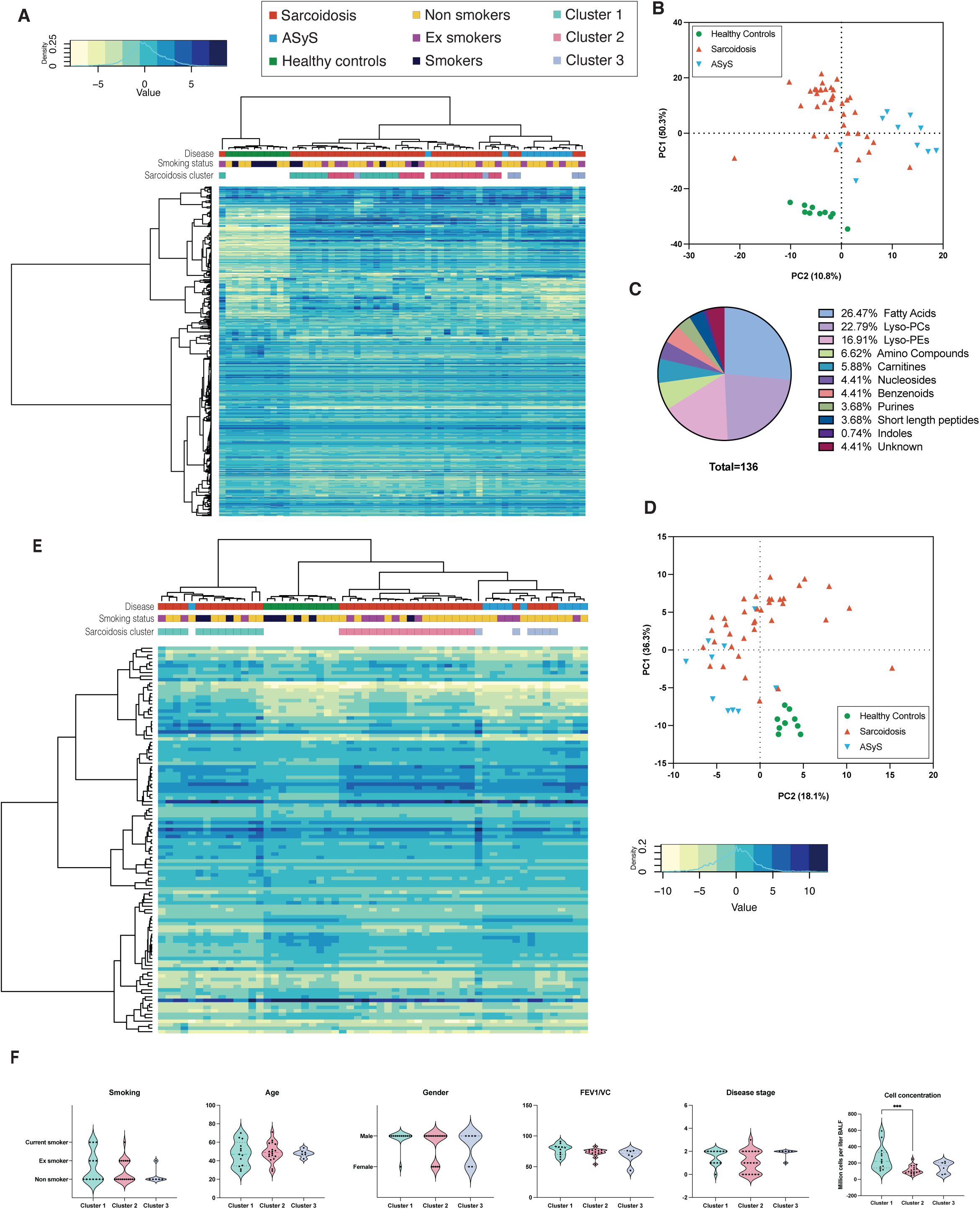
BALF EV samples originating from different diseases cluster together based on their metabolite cargos. A. Heatmap of all metabolite expressions across samples (Log2 transformed data, median-normalized, clustered by the ward algorithm). The analysis is based on all metabolites detected in the untargeted analysis, irrespective of their identification. B. Principal component analysis reveals a clustering of samples based on the disease they originate from when analyzing their total metabolite cargos. C. Pie chart representing the proportion of each class of metabolites or lipids detected in our targeted assay. D. Principal component analysis reveals a clustering of samples based on the disease they originate from when analyzing their metabolite cargos. E. Heatmap of metabolite expression across all samples (Log2 transformed data, median-normalized, clustered by the ward algorithm). F. Comparison of some clinical parameters smoking, age, gender, spirometry, disease stage, and cell concentration in the BALF between the 3 clusters of sarcoidosis patients. Significance is calculated by one-way ANOVA with Tukey’s multiple comparison test.

After annotated metabolomic analysis, 136 metabolites were detected in any of the samples. These metabolites stem from a variety of compound classes, with fatty acids, lysophosphatidylcholines (lyso-PCs) and lysophosphatidylethanolamines (lyso-PEs), and are strongly represented in the dataset (Figure 2C). After filtering, imputation, and normalization of the data, 111 individual metabolites remained for the differential expression analysis. The PCA revealed a clear separation of EV samples based on their disease of origin, with PC1 and PC2 explaining 36.3% and 18.1% of the total variance, respectively (Figure 2D). Additionally, a heatmap of metabolite content showed that the clinical status was the driving force of the clustering, separating healthy EVs from both Sarcoidosis and ASyS samples, and showing the specificity of EVs for their disease of origin (Figure 2E). Interestingly, both the non-annotated and the annotated heatmaps separated the sarcoidosis samples into three individual clusters. For the annotated data-based clusters, we compared the clusters with clinical information such as smoking, age, gender, spirometry, stage of the disease, and no significant effect was found due to these clinical parameters on the clustering. However, cell concentration in BALF samples was found significantly higher in cluster 1 compared to cluster 2 (Figure 2F). As smoking is an important risk factor in several lung disorders, the smoking status of the patients and healthy controls was also considered to see if it had any impact on the EVs’ metabolite cargo. However, no differences were detected, even when looking specifically at the effect of tobacco load in healthy controls (Supplementary Figure 1A) and sarcoidosis patients (Supplementary Figure 1B).

This exploratory overview of metabolites detected in EVs derived from patients and healthy donors shows that sarcoidosis and ASyS are the primary source of the variance in the metabolomic profile of EVs, separating healthy EVs from them, with some overlap between diseases.

### Individual metabolites are differentially expressed in the BALF EVs from patients with sarcoidosis and ASyS

EV samples originating from each disease were compared both between disease groups and to healthy controls, to investigate disease-specific metabolic signatures. The disease-specific signatures compared to healthy controls are represented in Figure 3A (Sarcoidosis) and 3B (ASyS). 87 and 59 metabolites were significantly different in sarcoidosis and ASyS patients, respectively, compared to healthy controls.

**Figure 3:**
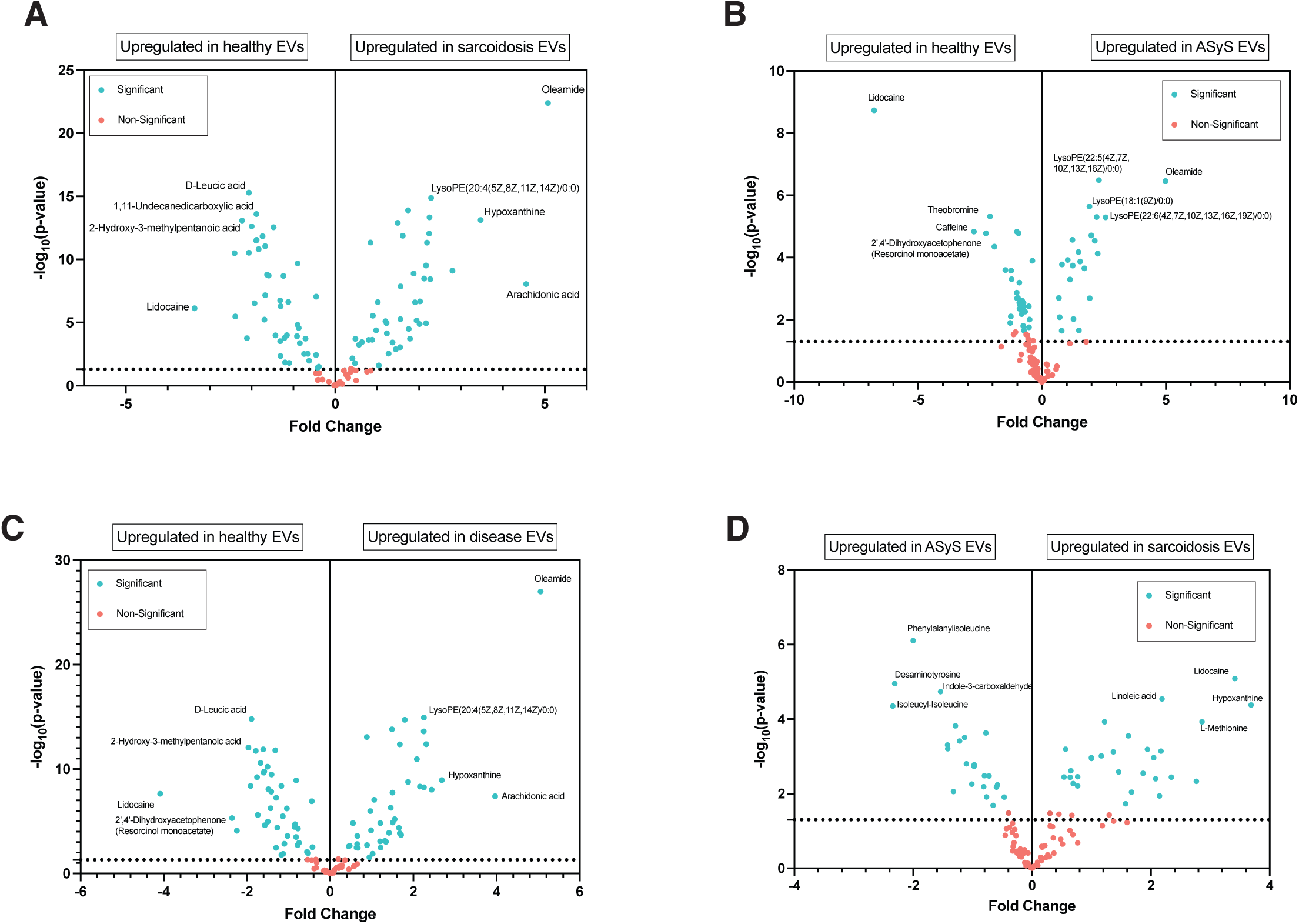
BALF EVs exhibit specific metabolites enriched or diminished in sarcoidosis or ASyS compared to healthy controls. A. Volcano plot representing the metabolites up-regulated (right) or down-regulated (left) in sarcoidosis EVs compared to healthy control EVs. B. Volcano plot representing the metabolites up-regulated (right) or down-regulated (left) in ASyS EVs compared to healthy control EVs. C. Volcano plot representing the metabolites up-regulated (right) or down-regulated (left) in EVs from patients with ASyS or sarcoidoisis, compared to healthy control EVs. D. Volcano plot representing the metabolites up-regulated (right) or down-regulated (left) in sarcoidosis EVs compared to ASyS EVs. For all volcano plots, the Log2 fold change is plotted against the p-value, and metabolites are highlighted in blue if the adjusted p-value (FDR) is lower than 0.05. The top 4 significant metabolites are labeled on each plot.

The EV composition of all patients (Both with sarcoidosis and ASyS) was also compared to that of healthy controls (Figure 3C), showing 86 metabolites significantly dysregulated. Additionally, the comparison of sarcoidosis and ASyS EVs (Figure 3D) shows a different metabolomic profile, with 56 metabolites differentially detected between the two groups, further highlighting the metabolomic differences between the diseases.

Overall, these findings demonstrate a distinct metabolic signature between each disease group and healthy individuals, indicating selective and distinct presentation of metabolites in the EVs of both patient groups.

### Differential expression of certain metabolites in EVs differentiates healthy individuals from those with diseases

When metabolites specific to healthy samples were investigated, we identified metabolites differentially expressed in only EVs derived from healthy EVs when compared to sarcoidosis and ASyS-derived EVs. Specifically, seven metabolites were found to be significantly overexpressed in healthy EVs (Figure 4A); two metabolites were found to be significantly downregulated in healthy EVs compared to both sarcoidosis and ASyS-derivedEVs (Figure 4B). Certain metabolites were also identified to be downregulated in sarcoidosis EVs compared to both healthy and ASySASys-derived EVs and lower in ASyS EVs than those from healthy (Figure-4C). Interestingly, only one specific metabolite, isoleucyl-isoleucine, was upregulated solely in ASyS EVs (Figure 4D). Among all these metabolites, specific lyso PCs and PEs were found to be differentially expressed between healthy and disease groups. While Lyso PEs were significantly overexpressed in Sarcoidosis and ASyS EVs compared to healthy EVs (Supplementary Figure 2A), no difference was identified in the expression level of Lyso PCs between the EVs from disease and healthy samples (Supplementary Figure 2B).

**Figure 4:**
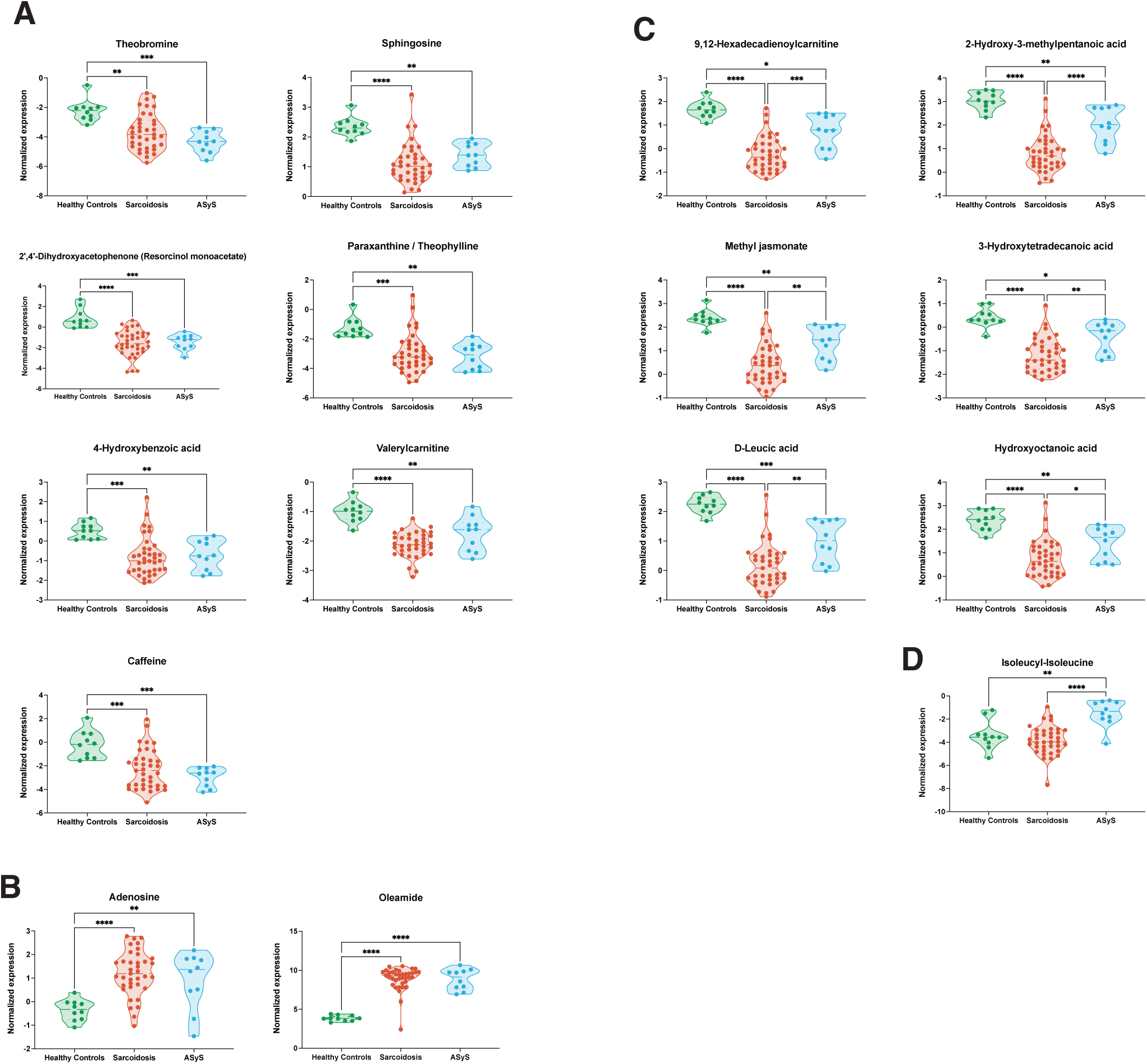
The metabolites are significantly different in EVs from both sarcoidosis and ASyS patients compared to healthy controls. A. Violin plot of all unique metabolites significantly downregulated in sarcoidosis and ASyS EVs compared to healthy controls. B. Violin plot of all unique metabolites significantly upregulated in sarcoidosis and ASyS EVs compared to healthy controls. C. Violin plot of all unique metabolites significantly downregulated in sarcoidosis and ASyS EVs compared to healthy controls, while being upregulated in AsyS compared to sarcoidosis. D. Violin plot of the unique metabolite significantly upregulated in ASyS EVs compared to those from both sarcoidosis patients and healthy controls. Significance is calculated by one-way ANOVA with Tukey’s multiple comparison test

These differential metabolite expression patterns in EVs from healthy individuals compared to those from sarcoidosis and ASyS, suggest that alterations in the metabolomic pathways are reflected in EVs and can help us identify candidate metabolites with potential to distinguish healthy individuals from both sarcoidosis and ASyS patients.

### Distinct metabolite signature is observed in sarcoidosis-derived EVs compared to ASyS and healthy EVs

In the next step, we explored a subset of metabolites that were significantly over- or under-represented in EVs from sarcoidosis patients compared to those from ASyS samples and healthy controls. 10 metabolites were found to be downregulated in only sarcoidosis EVs compared to both healthy and ASyS samples (Figure-5A). Additionally, 12 metabolites were identified as significantly upregulated in sarcoidosis EVs compared to ASyS and healthy EVs (Figure 5B). These metabolites were enriched in several metabolic pathways, including alpha linoleic acid metabolism, beta oxidation of very long chain fatty acids, betaine metabolism, and caffeine metabolism (Supplementary Figure 3). To validate the mass spectrometry (MS) analysis findings, ELISA was performed for Arachidonic Acid on EVs from sarcoidosis patients, yielding a strong correlation between the measured ELISA concentration and normalized MS expression (Figure 5C). As sarcoidosis can manifest in organs other than the lungs, we subsequently looked for the EV profile associated with extrapulmonary manifestations (EPM) and identified arachidonic acid among 6 other metabolites (docopentaenoic acid (22n-6), paraxanthine/theophylline, xanthine, phenylalanyliosleucine, resorcinol monoacetate, and niacinamide) that were significantly different in EVs from patients with or without EPMs with significant differences (Figure 5D).

**Figure 5:**
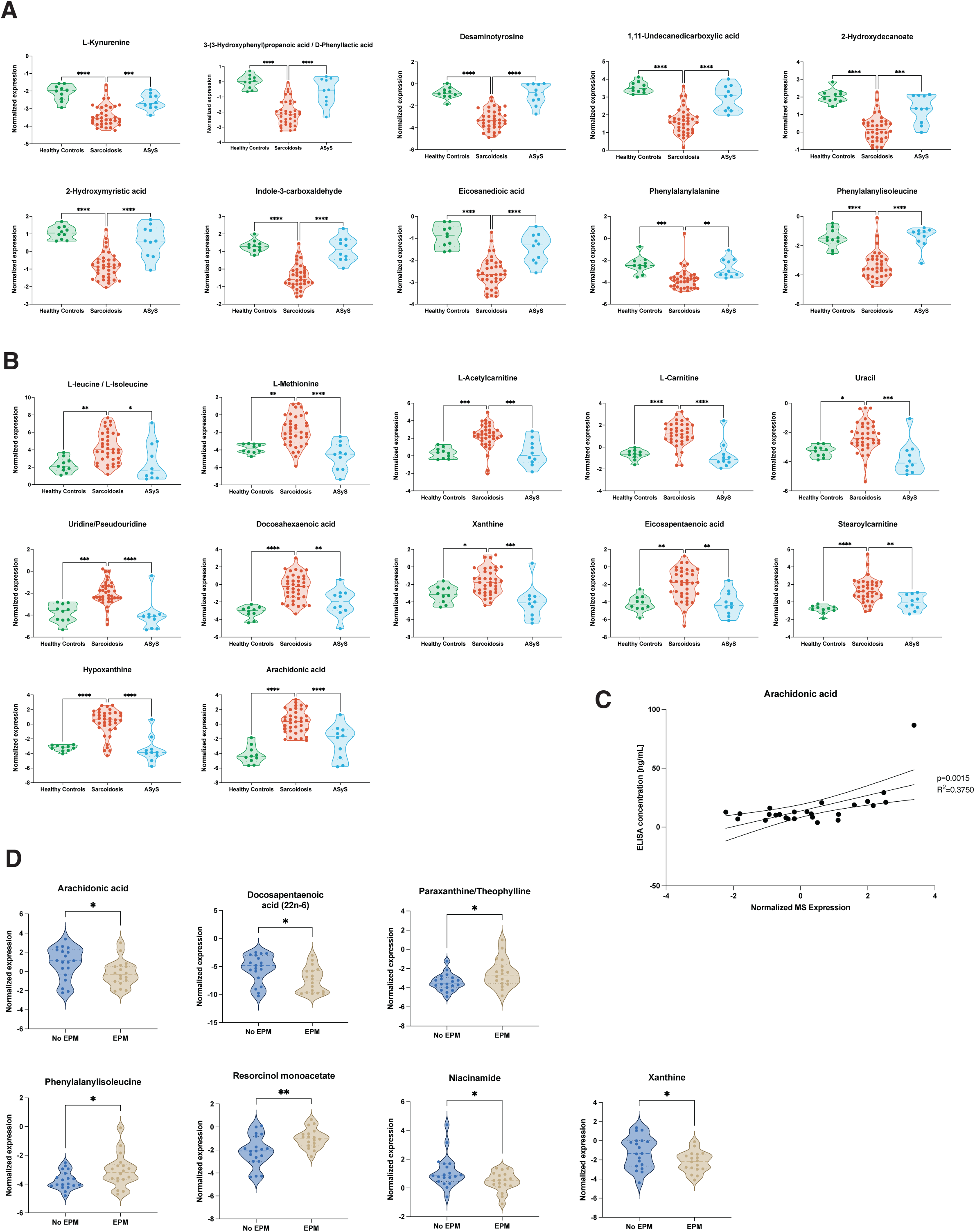
The specific metabolic signature in EVs from sarcoidosis and ASyS patients. A. Violin plot of all unique metabolites significantly downregulated in sarcoidosis EVs compared to those from both ASys patients and healthy controls. B. Violin plot of all unique metabolites significantly upregulated in sarcoidosis EVs compared to those from both ASys patients and healthy controls. C. Regression plot showing the correlation between the Arachidonic acid concentration detected by ELISA and arachidonic acid normalized MS expression in sarcoidosis EV samples. D. Representation of the 7 metabolites significantly associated with extrapulmonary manifestations (EPM). Significance in A-B and D are calculated by one-way ANOVA with Tukey’s multiple comparison test and t-test, respectively.

Altogether, these findings define a distinct arachidonic acid pathway signature in EVs from sarcoidosis patients, supported by independent ELISA validation, and further reveal subgroup heterogeneity based on EPM.

## Discussion

This study reports the first metabolomic characterization of EVs isolated from human BALF. It particularly focuses on the comparison of EVs from sarcoidosis and ASyS patients, alongside age-matched healthy controls. The comparison of sarcoidosis patients not only to healthy controls but also to another disorder that affects the lungs allows for the identification of novel disease-specific features, rather than merely studying the effects of an enhanced innate immune response, as previously discussed in the editorial to the first EV study in sarcoidosis (19, 21). This is further highlighted by the identification of uniquely upregulated metabolites in specific disorders. In addition, the untargeted analysis clearly demonstrates the clustering of samples based on the disease of origin (Figure 2A), looking at all metabolites present in EVs, even those not confidently identifiable based on their mass and retention time, which supports the idea that there is a specific metabolic identity for EVs, one that relates to the disorder of the human donor.

The similarities between EVs from patients with sarcoidosis and ASyS suggest one or several common disease mechanisms. This observation is also compatible with the notion that strong autoimmune components drive both sarcoidosis and ASyS, which is reflected in their respective metabolic signature of the EVs. Interestingly, the heatmap of metabolite abundance reveals the presence of three distinct clusters within the sarcoidosis samples, one of them overlapping with the majority of ASyS samples, emphasizing the close similarities between these two diseases in this respect. Of the clinical information available for the utilized study materials, only one parameter, total cell counts in BALF, showed a correlation to the clustering of sarcoidosis samples. Other data, such as smoking, spirometry, and age, did not show any striking differences between the sarcoidosis sub-clusters identified. It is interesting to speculate about factors segregating the patients into different subgroups. Such factors could be differences in disease triggers, disease progression, or prognosis, which are then reflected in the BALF EV metabolic composition. Understanding driving factors for subgrouping of patients in sarcoidosis might provide the necessary information to develop cures for this disease, and it would be interesting to follow the metabolic profile during the cause of disease and treatment. Furthermore, it would be interesting to compare EV data with previously suggested endotypes found by cell transcriptomic analyses (22).

Elevated levels of adenosine in serum were found in asthma and Chronic obstructive pulmonary disease (COPD) patients promoting fibrosis and disease progression (23, 24). Here, we found that adenosine is enriched in EVs from sarcoidosis and ASyS patients compared to healthy individuals. Our findings are consistent with reports highlighting the role of adenosine in chronic lung diseases. Furthermore, it supports our hypothesis that metabolomic alterations in sarcoidosis and ASyS patients can be captured in the metabolome of BALF EVs.

Caffeine and its three metabolites were found higher in healthy EVs than both disease EVs. These metabolites are methylxanthine alkaloids that are found in coffee, tea and chocolate indicating the potential effect of dietary habits on metabolites (PMID: 29514871). They are known for their bronchodilator properties (PMID: 26880379). In this study, we do not have information on dietary habits of healthy individuals and patients. However, our results correlate with the findings by Banoei et al, also showing decreased serum levels of theophylline and paraxanthine in sarcoidosis patients compared to healthy individuals (PMID: 39852350). These consistent findings from BALF EVs and serum metabolites might indicate an altered caffeine metabolism in sarcoidosis, possibly related to adenosine receptors. Caffeine as well as adenosine binds adenosine receptors, and altered receptor expression and/or competition could give an explanation to these findings (PMID: 21188209). Caffeine and its three metabolites were found to be higher in healthy EVs than EVs isolated from patients with either sarcoidosis or ASyS. These metabolites are methylxanthine alkaloids that are found in coffee, tea and chocolate indicating the potential effect of dietary habits on metabolites (25). They are known for their bronchodilator properties (26). Here, we do not have information on dietary habits of healthy individuals and patients. However, our results correlate with the findings by Banoei et al, also showing decreased serum levels of theophylline and paraxanthine in sarcoidosis patients compared to healthy individuals (27). These consistent findings from BALF EVs and serum metabolites might indicate an altered caffeine metabolism in sarcoidosis, possibly related to adenosine receptors. Caffeine as well as adenosine binds adenosine receptors, and altered receptor expression and/or competition could give an explanation to these findings (28).

Previously, we identified a distinct leukotriene pathway pattern in EVs from patients with sarcoidosis at the protein level (19, 29). In this study, we add to these observations that sarcoidosis patients have elevated levels of arachidonic acid, a precursor to the inflammatory mediators’ leukotriene family. In addition, omega-3 fatty acids Eicopentaeonic acid (EPA) and Docosahexaeonic acid (DHA), precursors to anti-inflammatory molecules, are enriched in sarcoidosis patients, which strengthens a potential link between EVs and disease-specific inflammatory processes. By focusing on the lipid mediators, this study further supports an alteration of the leukotriene pathway in sarcoidosis by showing not only differences in the leukotriene-forming enzymes but also in the lipid mediators upstream, downstream, or part of the leukotriene pathway. In support of this, Sjödin et al. have previously demonstrated that soluble epoxide hydrolase-derived lipid mediators were elevated in sarcoidosis patients (30), suggesting a disruption in arachidonic acid metabolism in sarcoidosis. Soluble epoxide hydrolases are involved in converting arachidonic acid into epoxyeicosatrienoic acids (EpETrEs) and dihydroxyeicosa-trienoic acids (DiHETrE), which have been shown to exert anti- and pro-inflammatory effects, respectively.

D-leucic acid (or 2-hydroxyisocaproic acid) is one of the compounds that has been found to be downregulated in EV’s from patients with ASyS and sarcoidosis, compared to those from healthy controls in our study. Sakko et al. (31) previously demonstrated that leucic acid exhibits anti-bacterial and anti-fungal properties, particularly against *Aspergillus*. Another anti-microbial compound effective against *Aspergillus*, D-Phenyllactic acid (32), has also been found to be downregulated in sarcoidosis EVs compared to both ASyS and healthy EVs. Interestingly, *Aspergillus Nidulans* has previously been proposed as a trigger of sarcoidosis (33). The downregulation of such fungicidal metabolites in the BALF EVs of patients could help seed a fungal infection in the lungs and promote disease initiation. Yet, further studies are needed to understand whether this information can be used for preventive measures or even early treatment.

Several studies have focused on systemic metabolic dysregulations associated with sarcoidosis (30, 34, 35), or linked them to clinical features of the disease (10). However, the contribution of EVs to these metabolic changes has not been studied yet. Some of our findings align with the existing literature on soluble metabolites, while others do not (particularly studies looking at plasma rather than BALF).

While Suzuki et al. previously showed that patients with sarcoidosis and multiple organ involvement had lower serum levels of soluble n-3 poly-unsaturated fatty acids (PUFAs) and a lower n-3/n-6 ratio than those with single organ involvement (10), we did not observe such correlations in our current study. Nevertheless, organ involvement is an uncertain clinical parameter. Indeed, our finding of increased Omega-3 in sarcoidosis patients compared to the other two disease groups supports an important role of PUFAs in sarcoidosis.

Common plasma metabolomic features in veterans with sarcoidosis were reported, whereas civilians with sarcoidosis showed distinctly different profiles (34). Indeed, in the current study, we also found strong differences in our (civilian) sarcoidosis, suggesting fundamentally distinct mechanisms underlying lung diseases induced by smoke or heavy particle exposure compared to those with a stronger genetic link.

Other studies showed altered phenylalanine metabolism in sarcoidosis patients either by elevated serum levels of the metabolites (35) and altered phneylalanine-to-tyrosine ratios reflecting immune activation (36). Lower levels of Phenylalanine containing dipeptides (phenylalanylalanine and phenylalanylisoluecine) and phenylalanine derived metabolite D-phenyllactic acid in sarcoidosis EVs support the alteration in phenylalanine metabolism indicating a potential role of EVs in lung inflammation.

The effect of exposure to cigarette smoke was investigated in the current study as well. However, the metabolic profiles of smokers were not statistically different compared to non-smokers. Therefore, we pooled these groups for further analysis. Although small groups of smokers were analyzed, which reduced the power of our analyses, the changes in quantities did not suggest large differences. Therefore, it is unlikely that increasing the sample size would result in substantial differences detected. Interestingly, the disease has a much stronger impact on the EV metabolome than smoking, even though smoking is known to alter lung cell composition and affect soluble metabolites in the lung. This may suggest that EV associated metabolites are more sensitive indicators of disease than soluble metabolites.

Dietary metabolites, including Omega-3, as discussed above, varied between patient groups, which could be explained by several factors. For instance, different disease groups may have different diets (consciously or not), which could be related to disease risks. Alternatively, disparities in dietary metabolism could also explain this finding, potentially due to an inborn metabolic defect that causes the disease itself. Another possibility is that the patients acquire an altered metabolism secondary to the disease, and the underlying types of inflammation. Lastly, variations in the gut permeability or the microbiome between patient groups might result in different food processing. It has been shown that sarcoidosis patients have a dysregulated microbiome (37), but whether this is causative or the effect of the disease is unclear. Differences in epithelial barrier integrity, including in the gut and/or lung, can cause different types of EV-metabolites to enter the pulmonary cavity. It has previously been shown that EVs with dietary protein antigens can be released from the gut epithelium and enter the blood stream (38) and it is possible that they are also detectable in the lungs, depending on epithelial integrity.

Additionally, as we analyzed EVs derived from BALF and not from serum or plasma, the comparison with literature and our understanding of the differences between BALF and blood EV metabolome is limited. Despite these limitations, this study provides a strong foundation for understanding the contribution of EVs to the metabolomic changes observed in different lung diseases, potentially opening the possibility for treatments with drugs interfering with lipid metabolism (39).

In conclusion, this study presents original evidence of metabolic differences in EVs isolated from the BALF of patients with sarcoidosis, ASyS and healthy controls, respectively. The distinct clustering of sarcoidosis and ASyS samples highlights differences that may prove useful for the development of novel diagnostic strategies and treatments.

## Acknowledgements

The expertise of the Swedish Metabolomics Center was greatly appreciated for the sample processing and preliminary analyses. This work was funded by the Swedish Heart-Lung Foundation (B.H. #20241216, A.L. #20210286, S.K. #20220140, S.K. #20220194, I.E.L. #20200379, G.G.A. #202000818, S.G. #20230528), the Swedish Research Council (A.L.: #2021-01527, S.K. #2022-00870, I.E.L. #2020-01378, S.G. #2022-01170, #2025-02722), the Karolinska Insitutet KID grant (SG), the Swedish Rheumatism Association (I.E.L.), King Gustaf V 80 Year Foundation (I.E.L.), Konung Gustaf V:s and Drottning Victorias Frimurarestiftelse (I.E.L.), the Stockholm County Council (S.K. FoUI-1001474), Regional agreement on medical training and clinical research (ALF) between Stockholm County Council and Karolinska Institutet (B.H. #202402457, S.K. #512827, S.K.#998890, I.E.L.), Region Stockholm (A.L.: #20180088), the Swedish Society for Respiratory Medicine (A.L.) and an unrestricted research grant from Astra Zeneca AB and Chiesi AB, Sweden (A.L.).

## Figure legends

**Supplementary Figure 1.**
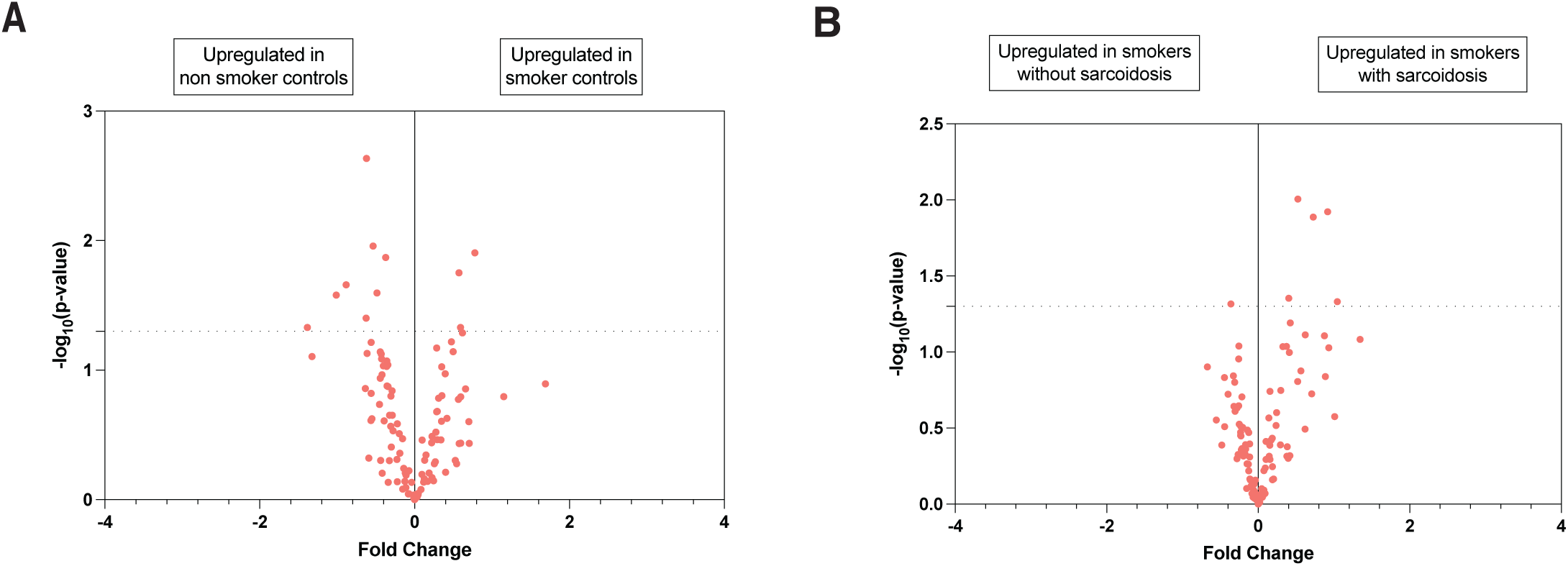
Volcano plots representing the metabolites A. Upregulated in smoker (right) and down-regulated in non-smoker healhy control EVs, B. Upregulated in EVs from smoker with sarcoidosis (right) and down-regulated in non-smoker sarcoidosis patients (left). For all volcano plots, the Log2 fold change is plotted against the p-value.

**Supplementary Figure 2:**
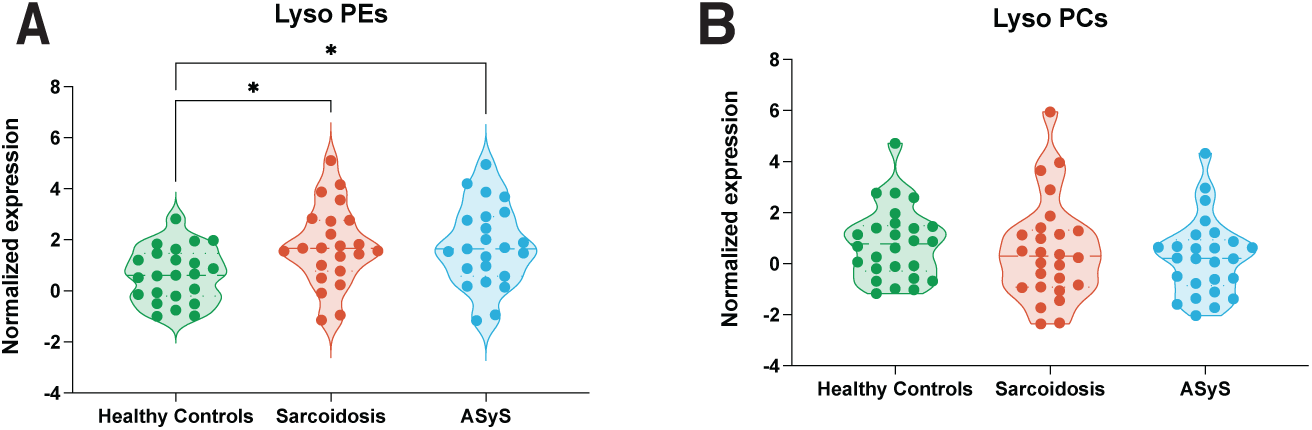
Violin plots comparing normalized expression of A. Lyso PEs and B. Lyso PCs in EVs from healthy controls, sarcoidosis and ASyS patients. Significance is calculated by one-way ANOVA with Tukey’s multiple comparison test.

**Supplementary Figure 3:**
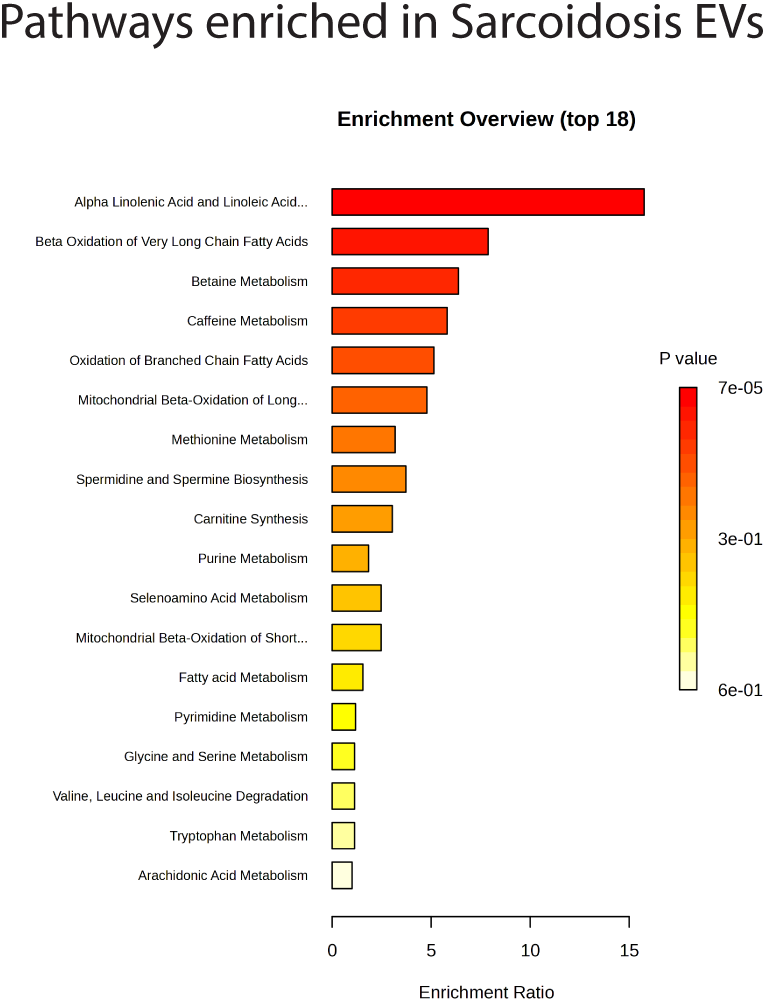
Enrichment analysis of the pathways differentially regulated in sarcoidosis EVs.

