## Supplementary table for "The metabolic profile of Extracellular Vesicles identifies and separates patients with Sarcoidosis and Anti-Synthetase Syndrome"

| Subject ID | Group | Smoking Status | Sex | Age | Stage | LS/Non-LS | Medication |
|---|---|---|---|---|---|---|---|
| I007 | Healthy | Non smoker | M | 54 | | | |
| I010 | Healthy | Non smoker | F | 41 | | | |
| I017 | Healthy | Non smoker | M | 46 | | | |
| I019 | Healthy | Non smoker | M | 52 | | | |
| I021 | Healthy | Non smoker | F | 70 | | | |
| R001 | Healthy | Smoker | M | 44 | | | |
| R011 | Healthy | Smoker | M | 41 | | | |
| R020 | Healthy | Smoker | F | 61 | | | |
| R022 | Healthy | Smoker | F | 58 | | | |
| R027 | Healthy | Smoker | F | 64 | | | |
| 1766 | Sarcoidosis | Ex smoker | M | 29 | I | LS | Inhalatory corticosteroid |
| 1645 | Sarcoidosis | Ex smoker | F | 42 | II | non-LS | Corticosteroids, MMF, TNF-inhibitors |
| 1649 | Sarcoidosis | Ex smoker | F | 31 | II | non-LS | Inhalatory corticosteroid |
| 1682 | Sarcoidosis | Ex smoker | F | 41 | II | LS | |
| 1684 | Sarcoidosis | Ex smoker | M | 45 | I | non-LS | |
| 1704 | Sarcoidosis | Ex smoker | M | 42 | II | non-LS | corticosteroids |
| 1725 | Sarcoidosis | Ex smoker | M | 70 | I | non-LS | |
| 1750 | Sarcoidosis | Ex smoker | M | 29 | I | non-LS | Inhalatory corticosteroid, Plaquenil |
| 1737 | Sarcoidosis | Ex smoker | M | 45 | II | non-LS | Inhalatory corticosteroids |
| 1747 | Sarcoidosis | Ex smoker | M | 71 | 0 | non-LS | corticosteroids |
| 1644 | Sarcoidosis | Non smoker | M | 54 | II | non-LS | corticosteroids |
| 1653 | Sarcoidosis | Non smoker | M | 47 | II | non-LS | corticosteroids, methotrexat |
| 1656 | Sarcoidosis | Non smoker | **F** | 58 | 0 | non-LS | corticosteroids, methotrexat |
| 1660 | Sarcoidosis | Non smoker | M | 35 | II | non-LS | |
| 1664 | Sarcoidosis | Non smoker | M | 53 | II | non-LS | corticosteroids |
| 1668 | Sarcoidosis | Non smoker | M | 47 | I | non-LS | topical corticosteroids (eye) |
| 1673 | Sarcoidosis | Non smoker | M | 64 | II | non-LS | |
| 1677 | Sarcoidosis | Non smoker | M | 62 | II | non-LS | |
| 1678 | Sarcoidosis | Non smoker | F | 60 | 0 | LS | |
| 1681 | Sarcoidosis | Non smoker | M | 46 | I | non-LS | |
| 1685 | Sarcoidosis | Non smoker | M | 47 | I | non-LS | |
| 1687 | Sarcoidosis | Non smoker | F | 48 | 0 | non-LS | corticosteroids |
| 1691 | Sarcoidosis | Non smoker | F | 50 | I | non-LS | |
| 1705 | Sarcoidosis | Non smoker | M | 59 | 0 | non-LS | |
| 1721 | Sarcoidosis | Non smoker | M | 47 | II | non-LS | corticosteroids, methotrexat |
| 1726 | Sarcoidosis | Non smoker | M | 46 | II | non-LS | |
| 1727 | Sarcoidosis | Non smoker | M | 50 | 0 | non-LS | methotrexate, corticosteroids |
| 1731 | Sarcoidosis | Non smoker | M | 52 | II | non-LS | |
| 1732 | Sarcoidosis | Non smoker | M | 34 | II | non-LS | methotrexate, corticosteroids |
| 1699 | Sarcoidosis | Non smoker | F | 50 | II | non-LS | |
| 1420 | Sarcoidosis | Non smoker | M | 49 | II | non-LS | |
| 1654 | Sarcoidosis | Smoker | M | 56 | 0 | non-LS | |
| 1688 | Sarcoidosis | Smoker | M | 65 | II | non-LS | |
| 1690 | Sarcoidosis | Smoker | M | 42 | I | non-LS | |
| 1735 | Sarcoidosis | Smoker | M | 31 | II | non-LS | Inhalatory budesonide, corticosteroid |
| 1648 | Sarcoidosis | Non smoker | M | 51 | I | non-LS | Inhalatory corticosteroid |
| 1672 | Sarcoidosis | Non smoker | M | 51 | III | non-LS | probably NSAID at the time for BALF |
| 511 | ASyS | Non smoker | F | 57 | | | Steroids, Cyclophosphamide, Mycophenolate Mofetil |
| 518 | ASyS | Non smoker | F | 55 | | | Steroids, Cyclophosphamide, Mycophenolate Mofetil |
| 670 | ASyS | Non smoker | M | 62 | | | na |
| 735 | ASyS | Ex smoker | M | 58 | | | Steroids, Methotrexate |
| 828/859 | ASyS | Ex smoker | M | 80 | | | Steroids, Cyclophosphamide, Mycophenolate Mofetil |
| 862 | ASyS | Non smoker | M | 56 | | | Steroids, Cyclophosphamide, Mycophenolate Mofetil, Rituximab |
| 934 | ASyS | Ex smoker | F | 47 | | | Steroids, Cyclophosphamide, Mycophenolate Mofetil, Rituximab |
| 1615 | ASyS | Non smoker | M | 43 | | | Steroids, Mycophenolate mofetil, Rituximab |
| 1743 | ASyS | Non smoker | F | 33 | | | Steroids, Mycophenolate mofetil, Rituximab |
| 1794 | ASyS | Non smoker | F | 65 | | | Steroids, Methotrexate, Mycophenolate mofetil |
